# The novel bacterial effector protein *Cb*EPF1 mediates ER-LD membrane contacts to regulate host lipid droplet metabolism

**DOI:** 10.1101/2023.12.11.571031

**Authors:** Rajendra Kumar Angara, Arif Sadi, Stacey D. Gilk

**Affiliations:** Department of Pathology and Microbiology, University of Nebraska Medical Center, Omaha, Nebraska, USA

**Keywords:** Inter-organelle contacts, FFAT motif, Lipid droplets, Molecular tethers, Effector proteins, *Coxiella burnetii*

## Abstract

Effective intracellular communication between cellular organelles is pivotal for maintaining cellular homeostasis. Tether proteins, which are responsible for establishing membrane contact sites between cell organelles, enable direct communication between organelles and ultimately influence organelle function and host cell homeostasis. While recent research has identified tether proteins in several bacterial pathogens, their functions have predominantly been associated with mediating inter-organelle communication specifically between the bacteria containing vacuole (BCV) and the host endoplasmic reticulum (ER). However, this study reveals a novel bacterial effector protein, *Cb*EPF1, which acts as a molecular tether beyond the confines of the BCV and facilitates interactions between host cell organelles. *Coxiella burnetii*, an obligate intracellular bacterial pathogen, encodes the FFAT motif-containing protein *Cb*EPF1 which localizes to host lipid droplets (LDs). *Cb*EPF1 establishes inter-organelle contact sites between host LDs and the ER through its interactions with VAP family proteins. Intriguingly, *Cb*EPF1 modulates growth of host LDs in a FFAT motif-dependent manner. These findings highlight the potential for bacterial effector proteins to impact host cellular homeostasis by manipulating inter-organelle communication beyond conventional BCVs.

## INTRODUCTION

Inter-organelle communication, the dynamic exchange of biomolecules, ions, and lipids between cellular organelles, is integral to maintains cellular homeostasis. In the past decade, extensive research has unveiled inter-organelle membrane contact sites where key organelles, including the endoplasmic reticulum (ER), mitochondria, Golgi apparatus, peroxisomes, and lipid droplets (LDs), engage in direct communication (Elbaz and Schuldiner, 2011; Helle et al, 2013; Gatta and Levine, 2017; Wu et al, 2018; Scorrano et al, 2019). These inter-organelle contact sites are orchestrated by unique proteins with distinct motifs and domains that act as molecular tethers between organelles and facilitate biomolecule exchange (Eisenberg-Bord et al, 2016; Gatta and Levine, 2017; Silva et al, 2020). These sites serve as hubs for lipid transfer, calcium signaling, and membrane trafficking, and profoundly influence cellular processes such as lipid metabolism, signal transduction, and organelle biogenesis (Prinz et al, 2014; Wu et al, 2018; Bohnert et al, 2020; Prinz et al, 2020; Silva et al, 2020).

Pathogens employ intricate strategies to co-opt host cell machinery and manipulate cellular metabolism to ensure their survival and replication. Recent studies have revealed the pathogen’s ability to modify host inter-organelle communication (Kors et al, 2022; Jiang et al, 2022). Upon host cell entry, certain obligate intracellular bacteria adopt a unique endocytic membrane-bound vacuole, referred to as the Bacteria-Containing Vacuole (BCV), to evade the host immune response and support bacterial replication (Santos and Enninga, 2016; Petit and Lebreton, 2022). While the BCV shields bacteria, it restricts access to host cellular machinery and nutrients. Consequently, various types of secretion systems have evolved in different bacteria to facilitate delivery of bacterial effector proteins into host cells (Mitchell et al., 2017). Bacterial effectors localize to various host cell organelles, using molecular mimicry to manipulate host metabolism and promote bacterial survival. Recent studies have unveiled that, in addition to adopting eukaryotic catalytic and receptor domains as a form of molecular mimicry, bacterial effector proteins can also contain features of eukaryotic tether proteins and establish inter-organelle contact sites between the host ER and BCV membranes (Murray et al., 2017; Vormittag et al., 2023). Remarkably, although several bacterial effector proteins localize to host cell organelles outside the context of the BCV, none have been identified to mediate inter-organelle contact sites.

The ER serves as a central hub at the heart of inter-organelle communication (Phillips and Voeltz, 2016; Wu et al, 2018; Almeida et al, 2020). The majority of ER-driven inter-organelle communication is facilitated by FFAT (two phenylalanines in an acidic tract) motifs found in a diverse family of proteins (Murphy and Levine, 2016). The FFAT motif-containing proteins act as molecular tethers between ER-resident VAPs (VAMP-associated proteins) and various organelles (Neefjes and Cabukusta, 2021). Besides their role as tethers, FFAT motif-containing proteins can also function as lipid or ion transporters. While considered a eukaryotic motif, FFAT motifs are also found in secreted bacterial effector proteins which mediate inter-organelle contact sites between the ER and BCVs (Murray et al, 2017). Given that a substantial portion of intracellular bacteria effector proteins remain uncharacterized, it is plausible that many unidentified FFAT motif-containing effector proteins localize to host cell organelles and facilitate host inter-organelle communication.

Building upon existing knowledge of inter-organelle communication, our study investigates a unique *Coxiella burnetii* effector protein that participates in inter-organelle communication beyond BCVs. The *C. burnetii* effector protein containing FFAT motifs, termed *Cb*EPF1, establishes inter-organelle contact sites between the host ER and LDs. *Cb*EPF1 features two FFAT motifs, each demonstrating preferential interaction with VAPs in the ER. *Cb*EPF1 mediates ER-LD inter-organelle contact sites to promote development of larger LDs in an FFAT motif-dependent manner, offering insights into how *C. burnetii* reprograms host lipid metabolism.

## RESULTS

### *Cb*EPF1 localizes to host ER and LDs

To identify effector proteins that may establish inter-organelle contact sites beyond BCVs, we conducted a bioinformatic screening of 127 predicted *C. burnetii* Type 4B Secretion System (T4BSS) effector proteins (Chen et al, 2010 and Bi et al, 2013) for FFAT motifs using FIMO (Find Individual Motif Occurrences) (Grant et al, 2011). From this screen, CBU1370 contained the highest probability with two FFAT motif sequences, both bearing strong similarity to the consensus sequences and preceded by acidic amino acids (**Fig. 1A**)(Neefjes and Cabukusta, 2021). Previous studies confirmed CBU1370 as T4BSS effector protein (Lifshitz et al, 2014). Beyond the predicted FFAT motifs, the CBU1370 protein sequence contains a putative amphipathic helix at the C-terminal end (from 262–279 amino acids) but has no other identifiable conserved domains or motifs. Accordingly, we designated CBU1370 as *<u>C</u>oxiella <u>b</u>urnetii* <u>E</u>ffector <u>P</u>rotein with <u>F</u>FAT motifs <u>1</u> (*Cb*EPF1).

**Figure 1:**
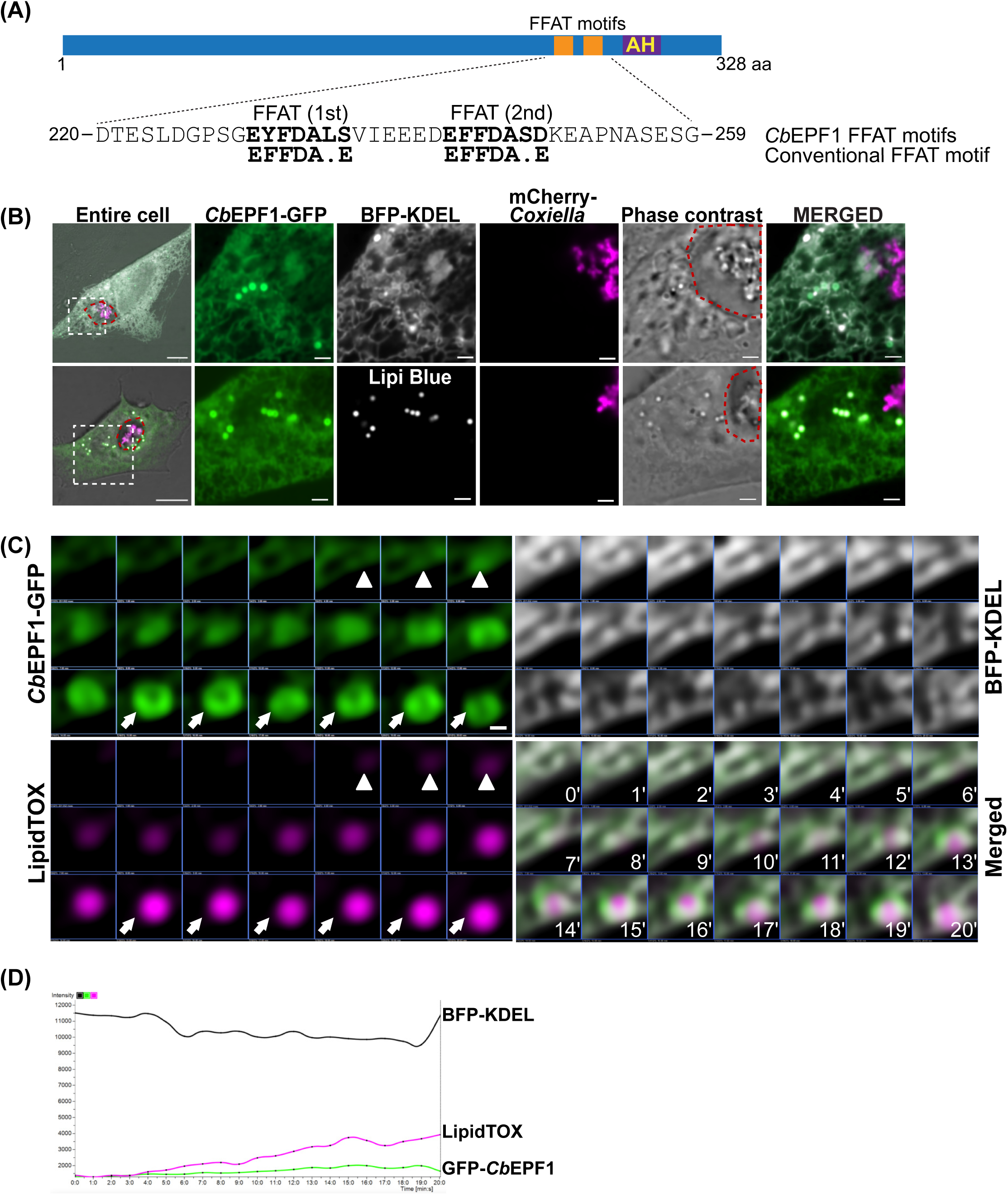
*Cb*EPF1-GFP localizes to host ER and LDs. (A) Two putative FFAT motifs were identified in the C-terminal region of *Cb*EPF1 protein. The sequence and position of the FFAT motifs are shown, with amino acid numbers indicated. Conventional FFAT motif sequence is shown below predicted *Cb*EPF1 FFAT motifs. AH represents a predicted site of amphipathic helix. (B) Live cell microscopy shows that ectopically expressed *Cb*EPF1-GFP localized to the host ER and LDs in mCherry-*Coxiella* infected HeLa cells. The ER was labeled with BFP-KDEL and LDs were labeled with LipidTOX-red. The *Coxiella* Containing Vacuole (CCV) membrane is outlined in red in the phase image. Scale bar 10 µm (overview) and 2 µm (magnified). (C) *Cb*EPF1-GFP associated with LD biogenesis in the ER. Arrowheads indicate early-stage LD biogenesis sites (t = 4-6 mins) and arrows mark larger LDs with *Cb*EPF1-GFP localized on the entire LD surface (t >13 mins). Images were acquired every 1 min (t = 0-20 min) by spinning disc confocal microscopy. Scale bar 0.2 µm. (D) Intensity profile of BFP-KDEL, *Cb*EPF1-GFP, and LipidTOX-red from Fig. 1C during LD biogenesis in the ER from t=0-20 min. From 4 min time point, the increase in LipidTOX-red intensity indicates emergence and growth of LD.

To understand the functional significance of *Cb*EPF1 in host cells, we ectopically expressed *Cb*EPF1-GFP in mCherry-*C. burnetii* infected epithelial cells. *Cb*EPF1-GFP predominantly localizes to the host ER with enrichment at distinct loci within the host cytosol (**Fig. 1B top panel**). Using co-localization with several organelle and vesicular markers, we determined that *Cb*EPF1-GFP also colocalizes with host LDs (**Fig. 1B bottom panel**). As LD biogenesis occurs at the ER (Kassan et al, 2013; Choudhary et al. 2015), we next used live cell imaging to examine the spatial and temporal localization of *Cb*EPF1 during LD biogenesis. We found that *Cb*EPF1-GFP initially localizes to ER sites of LD biogenesis and gradually enriches on the LD surface as LDs grow (**Fig. 1C and 1D**). Overall, these results demonstrate that *Cb*EPF1 dynamically distributes between the ER, LD biogenesis sites and the LD surface.

### *Cb*EPF1 relocates from ER to the LD surface and induces inter-organelle contact sites between the host ER and LDs

To gain deeper insight into *Cb*EPF1 localization dynamics at the ER and LDs, we induced LD synthesis by supplementing the media with oleic acid (OA). As LD number increased over time, we observed a corresponding decrease in *Cb*EPF1-GFP associated with the ER (**Fig. 2A**). In tandem, there was noticeable enrichment of *Cb*EPF1-GFP on host LD surfaces. This suggests a dynamic translocation of *Cb*EPF1-GFP from the ER to the LD surface, a process that coincides with the increase in LD size (**Fig. 2A**). Several eukaryotic proteins implicated in LD biogenesis or growth exhibit similar translocation pattern from the ER to LD surface as the LD grow to larger size (Wilfiling et al, 2013; Xu et al, 2012; Chung et al, 2019; Li et al, 2019). While larger LDs often detach from the ER and are released into the cytosol, they also often maintain contact with the ER by other mechanisms (Hugenroth and Bohnert, 2020). In our studies, cells expressing *Cb*EPF1-GFP presented a distinctive pattern, with the ER enveloping the *Cb*EPF1-GFP-localized LDs. The ER exhibited notably increased and extended association around *Cb*EPF1-GFP-localized LDs (**Fig. 2B and C**). In contrast, in control GFP-expressing cells the ER forms a reticulate network with only limited association with LDs (**Fig. 2B and 2C**). Remarkably, the ER wrapping around *Cb*EPF1-localized LDs was not always continuous but was distinctly confined to regions where *Cb*EPF1 localized on the LDs (**Fig. 2B and C**). This indicates that *Cb*EPF1 initiates and mediates contact sites between the ER and *Cb*EPF1-localized LDs, emphasizing its significance in orchestrating inter-organelle interactions.

**Figure 2:**
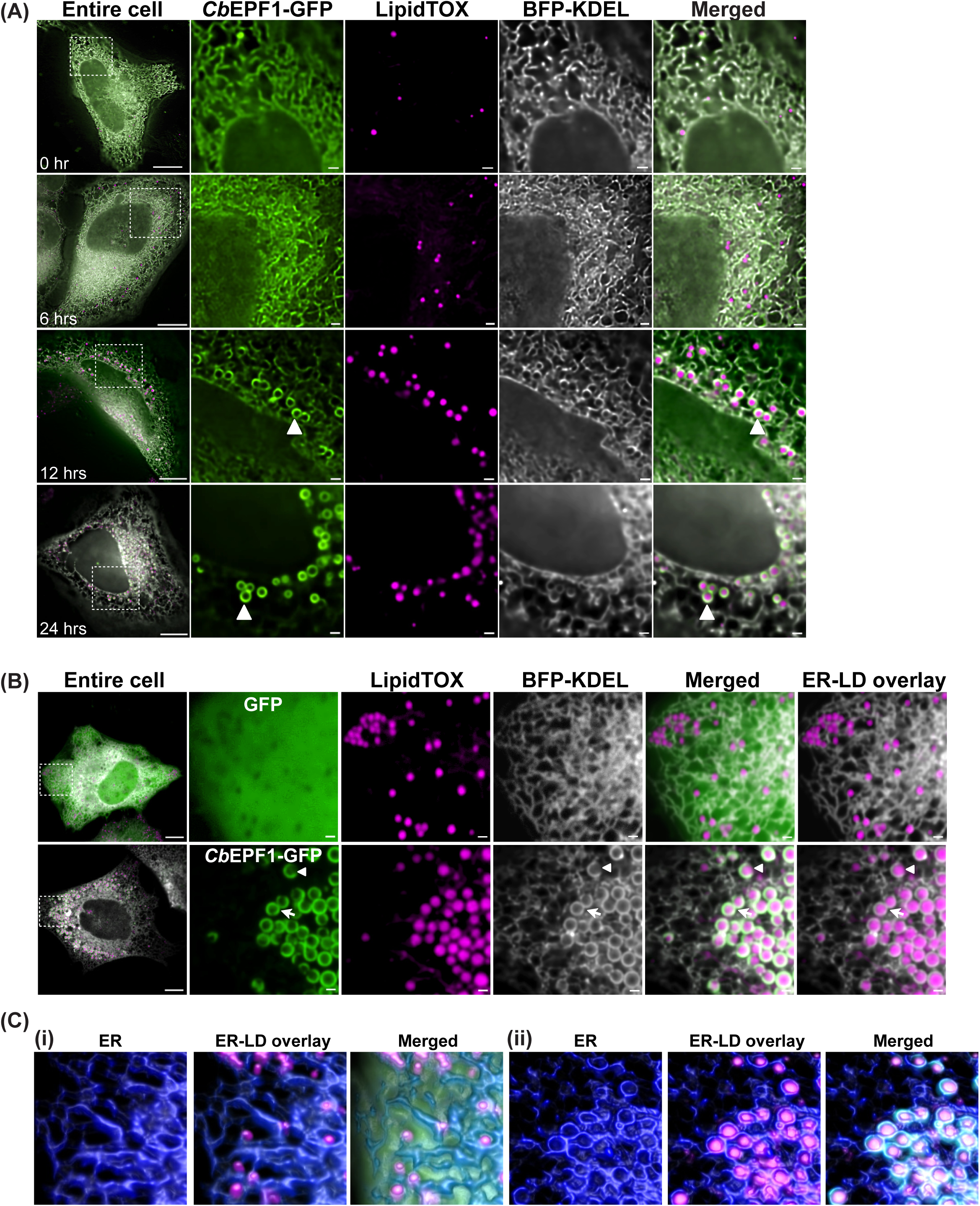
*Cb*EPF1 associates with host LDs and induces inter-organelle contact sites between the host ER and LDs. (A) HeLa cells expressing *Cb*EPF1-GFP and BFP-KDEL were treated with OA (30 µM) and imaged every 6 hrs by live cell spinning disc microscopy. Representative deconvoluted images show *Cb*EPF1-GFP relocates from the ER to the LD surface as LDs grow (arrowhead). Scale bar 10 µm (overview) and 1 µm (magnified). (B) HeLa cells expressing GFP or *Cb*EPF1-GFP and BFP-KDEL were treated with OA (100 µM, overnight) to induce LD biogenesis. LDs were visualized with LipidTOX red. ER shows limited association with LDs in GFP expressing cells (top panel). *Cb*EPF1-GFP expression induces extended contacts between ER and LDs (bottom panel). Arrows show ER wrapping around *Cb*EPF1 localized LDs. Arrow heads show absence of ER interaction at the regions of LDs where *Cb*EPF1-GFP localization is absent. Scale bar 10 µm (overview) and 1 µm (magnified). (C) Representative image of 3D rendering of a Z-stack image illustrating the association between LDs (magenta) and ER (blue). In cells expressing GFP alone (i), the ER exhibits minor contacts with LDs. In contrast, cells expressing *Cb*EPF1-GFP (ii) display extended ER-LD contacts, with ER-LD interactions specifically localized to regions where *Cb*EPF1-GFP (green) is present on LDs.

### *Cb*EPF1 contains two FFAT motifs and interacts with ER VAP proteins

To understand the factors responsible for *Cb*EPF1-mediated ER-LD contact sites, we focused on the *Cb*EPF1 FFAT motifs. VAPs in the ER membrane form heterodimeric complexes and establish inter-organelle contact sites between the ER and various organelles (**Fig. 3A**) (Murphy et al, 2016). VAPs, specifically VAPA, VAPB, and MOSPD2, form inter-organelle contact sites through the VAP MSP domain which binds the FFAT motif on partner proteins found on other organelles (Cabukusta et al, 2020). Consequently, proteins harboring FFAT motifs typically act as molecular tethers at inter-organelle contact sites. To evaluate whether *Cb*EPF1 binding to VAPs is responsible for *Cb*EPF1-mediated ER-LD contacts, we tested *Cb*EPF1-VAPB interactions using an *in vitro* Bacterial Adenylate Cyclase-based Two Hybrid assay (BACTH). In the BACTH assay, *Cb*EPF1 interacted with wildtype VAPB but not with a mutant VAPB containing a mutated MSP domain (VAPB-MSPmt), the VAP domain critical for interaction with FFAT motif containing proteins (**Fig. 3C**). To determine whether the *Cb*EPF1 FFAT motifs are required for binding to VAPB, we generated mutations in the two predicted *Cb*EPF1 FFAT motifs. While amino acids substitutions at most positions within the FFAT motif do not affect function, the phenylalanine or tyrosine residue at position 2 is indispensable for effective VAP binding (Murphy et al, 2016). Therefore, to test the functional importance of the putative *Cb*EPF1 FFAT motifs, we introduced alanine substitutions in position 2 of the first FFAT motif (Y231A, referred as F1mt), the second FFAT motif (F244A, referred as F2 mt), or both FFAT motifs (Y231A-F244A, referred to as YF/AA or F3mt) (**Fig. 3B**). In the BACTH assay, wildtype VAPB (VAPB-Wt) interacted with *Cb*EPF1-F1mt and *Cb*EPF1-F2mt but not with *Cb*EPF1-F3mt (**Fig. 3C**). These results indicate that *Cb*EPF1 interacts with VAPB through the FFAT and MSP domains. Further, while both *Cb*EPF1 FFAT motifs can bind VAPB, at least one functional FFAT motif is required and sufficient for *Cb*EPF1-VAPB binding. While no additional *C. burnetii* effector or eukaryotic factors are required for *Cb*EPF1-VAPB interactions *in vitro*, we cannot exclude the association/involvement of other proteins at the *Cb*EPF1–VAPB molecular tether.

**Figure 3:**
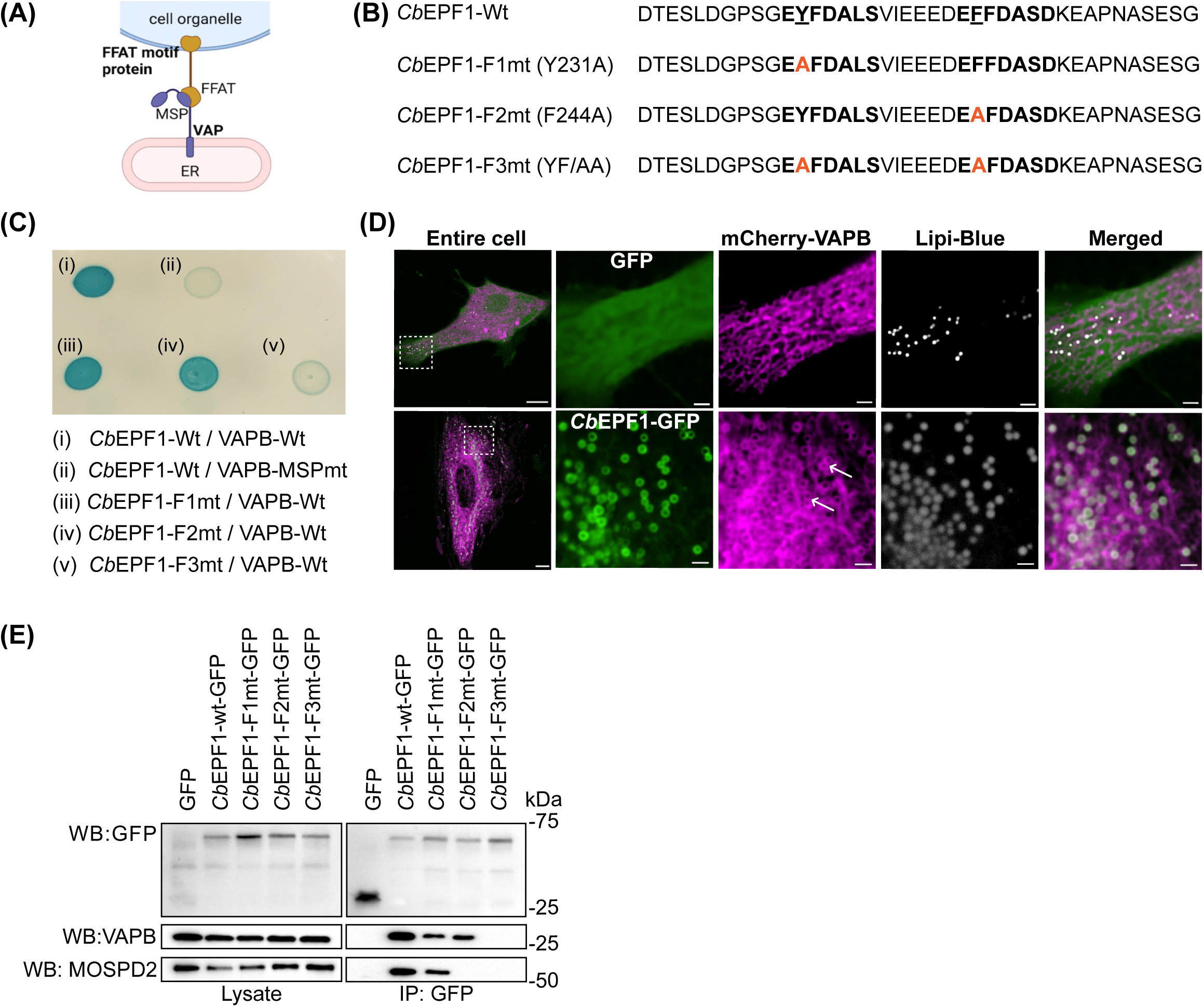
*Cb*EPF1 FFAT motifs interact with the VAP MSP domain. (A) Schematic representation of the FFAT motif containing protein interaction with the ER protein VAP to form inter-organelle contact sites between ER and LDs. (B) *Cb*EPF1 amino acid sequence contains two FFAT motifs. Number indicates amino acid position; essential position (2^nd^ residue) in FFAT motif is underlined. Mutations in FFAT motif(s) are highlighted in orange. (C) Bacterial adenylate cyclase-based two hybrid assay (BACTH) shows *Cb*EPF1 interacts with VAPB in an MSP-FFAT motif dependent manner. (D) HeLa cells expressing mCherry-VAPB along with GFP or *Cb*EPF1-GFP were treated with OA (100 µM, overnight) and LDs were stained with Lipi Blue. The white arrow shows mCherry-VAPB as ring like structure around *Cb*EPF1-GFP localized LDs. Scale bars: 10 µm (overview) and 2 µm (magnified). (E) Immunoprecipitation of GFP and *Cb*EPF1-GFP (WT, F1mt, F2mt, and F3mt) from lysates of HEK293 induced with OA (100 µM, overnight). WB represents western blot analysis using respective primary antibody.

### *Cb*EPF1 FFAT motifs show preferential interaction among VAP family proteins

We next sought to validate *Cb*EPF1 interaction with VAPs in mammalian cells. First, we ectopically expressed mCherry-VAPB along with GFP or *Cb*EPF1-GFP in HeLa cells and examined VAPB co-localization with respect to *Cb*EPF1 and LDs. In GFP-expressing cells, mCherry-VAPB exhibited a reticulate pattern and less interaction with LDs (**Fig. 3D, top panel**). In contrast, *Cb*EPF1-GFP expressing cells show mCherry-VAPB as ring-like structure around *Cb*EPF1-GFP positive LDs (**Fig. 3D, bottom panel**). Our observations in GFP control cells support previous studies that noted a lack of VAPB association with LDs (Zouiouich et al, 2022). Hence, we hypothesize that VAPB localization around LDs is due to *Cb*EPF1-GFP.

To further evaluate *Cb*EPF1 interaction with VAPs in mammalian cells, we expressed *Cb*EPF1-GFP or its FFAT mutant variants in HEK293T cells and induced LD formation with OA prior to immunoprecipitation. In western blot analysis, VAPB co-immunoprecipitated with *Cb*EPF1-GFP, *Cb*EPF1-F1mt-GFP and *Cb*EPF1-F2mt-GFP but not with *Cb*EPF1-F3mt-GFP or GFP (**Fig. 3E**). This suggests VAPB interacts with both *Cb*EPF1 FFAT motifs. Interestingly, another VAP family protein, MOSPD2, coimmunoprecipitated with *Cb*EPF1-GFP and *Cb*EPF1-F1mt-GFP but not with *Cb*EPF1-F2mt-GFP and *Cb*EPF1-F3mt-GFP. This suggests that unlike VAPB, MOSPD2 interacts with only the second *Cb*EPF1 FFAT motif (**Fig. 3E**). These data therefore demonstrate that the two FFAT motifs in *Cb*EPF1 display preferential interaction for VAP family proteins.

### *Cb*EPF1 FFAT motifs mediate ER-LD contacts

As we demonstrated that the two *Cb*EPF1 FFAT motifs and the VAP MSP domain are responsible for *Cb*EPF1–VAP interaction, we next analyzed whether the *Cb*EPF1 FFAT motifs mediate ER-LD contact sites. Using ectopic expression of GFP fusion proteins, we studied the role of each FFAT motif in *Cb*EPF1-mediated ER-LD contact sites, based on association of the ER with LDs. Mutations in the *Cb*EPF1 FFAT motif(s) did not influence *Cb*EPF1-GFP localization to either the host ER or LDs (**Supplementary Fig. 1**). However, only *Cb*EPF1-F1mt-GFP and *Cb*EPF1-F2mt-GFP proteins induce ER-LD contacts similar to *Cb*EPF1-Wt, whereas *Cb*EPF1-F3mt-GFP failed to induce ER-LD contacts (**Fig. 4A and 4C**). These results suggest that at least one functional FFAT motif is necessary and sufficient for *Cb*EPF1 to mediate ER-LD contact sites and rearrange the ER around LDs. Interestingly, *Cb*EPF1-F3-mt-GFP expression in cells led to LD clustering and exclusion of the ER in the vicinity of LD clusters (**Fig. 4B and 4C**). Since the *Cb*EPF1 FFAT motifs are essential for interaction with VAP proteins in the ER, the absence of functional FFAT motifs in *Cb*EPF1-F3mt-GFP disrupts the interactions between LDs and the ER. The observed clustering of *Cb*EPF1-F3mt-GFP localized LDs is likely due to the absence of LD interaction with the ER which may facilitate LD-LD interactions, potentially through interactions with unidentified proteins on the LD surface.

**Figure 4:**
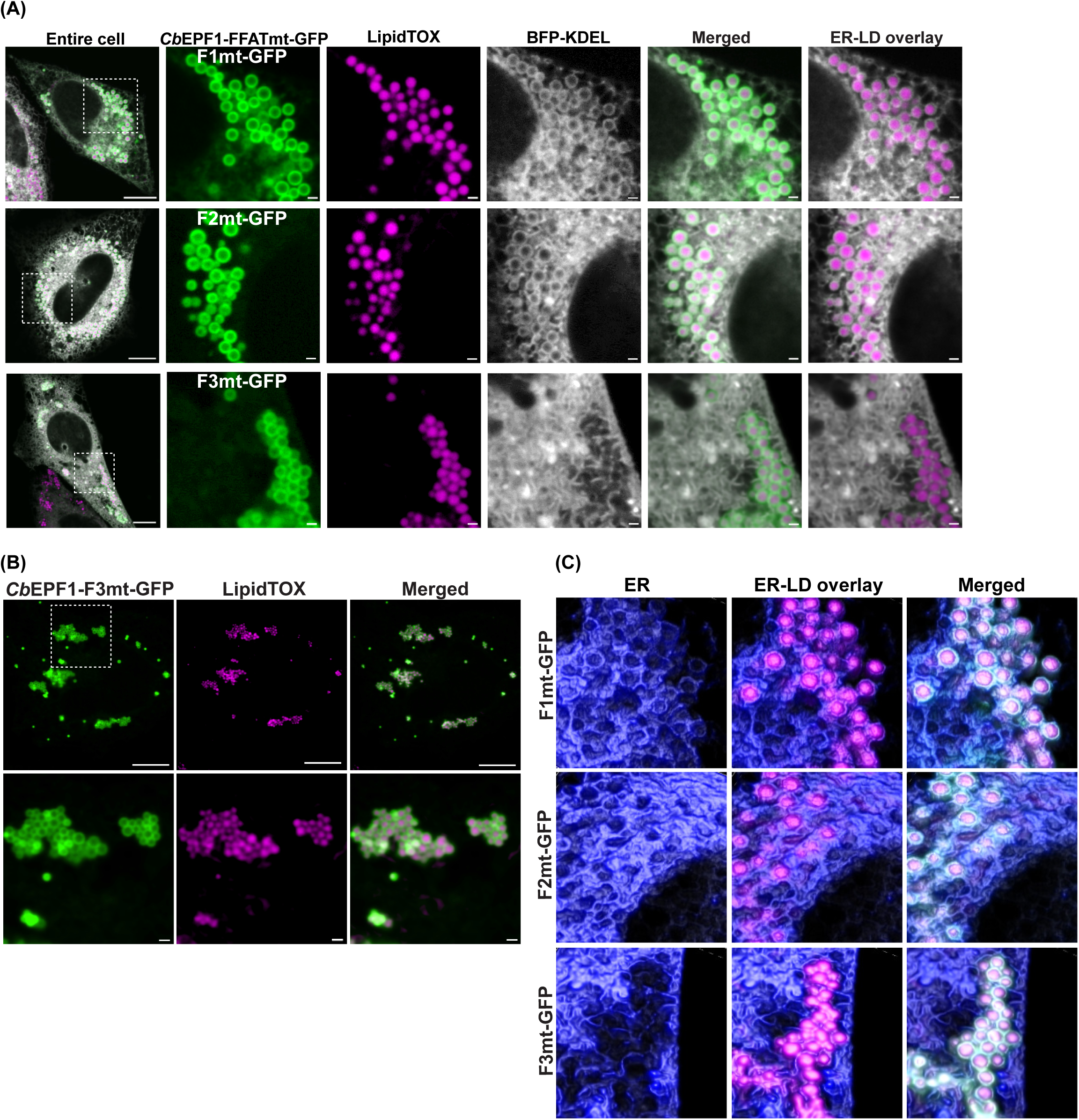
Functional FFAT motifs are required for *Cb*EPF1-induced ER-LDs contact sites. (A) *Cb*EPF1-F1mt-GFP and *Cb*EPF1-F2mt-GFP induce ER-LD contact sites while *Cb*EPF1-F3mt-GFP failed to induce ER-LD contact sites. Scale bar 10 µm (overview) and 1 µm (magnified). (B) *Cb*EPF1-F3mt-GFP expression in cells causes clustering of LDs. Scale bar 10 µm (overview) and 1 µm (magnified). (C) Representative image of 3D rendering of a Z-stack image illustrating the association between LDs (magenta) and ER (blue) in *Cb*EPF1-FFAT mutant-GFP expressing cells. *Cb*EPF1-F1mt-GFP or *Cb*EPF1-F2mt-GFP expressing cells exhibited ER-LD contacts, while the *Cb*EPF1-F3mt-GFP expressing cells showed exclusion of ER around LD clusters.

### *Cb*EPF1 regulates host LD metabolism in a FFAT-dependent manner

Next, we sought to evaluate whether *Cb*EPF1 impacts host LD metabolism by measuring LD numbers in cells expressing *Cb*EPF1-GFP or the *Cb*EPF1-GFP FFAT mutants. Compared to cells expressing GFP alone, cells expressing *Cb*EPF1-GFP or *Cb*EPF1-FFAT mutant GFP showed significantly high number of LDs (**Fig. 5A**), suggesting *Cb*EPF1 expression impacts LD number independent of the FFAT motif.

**Figure 5:**
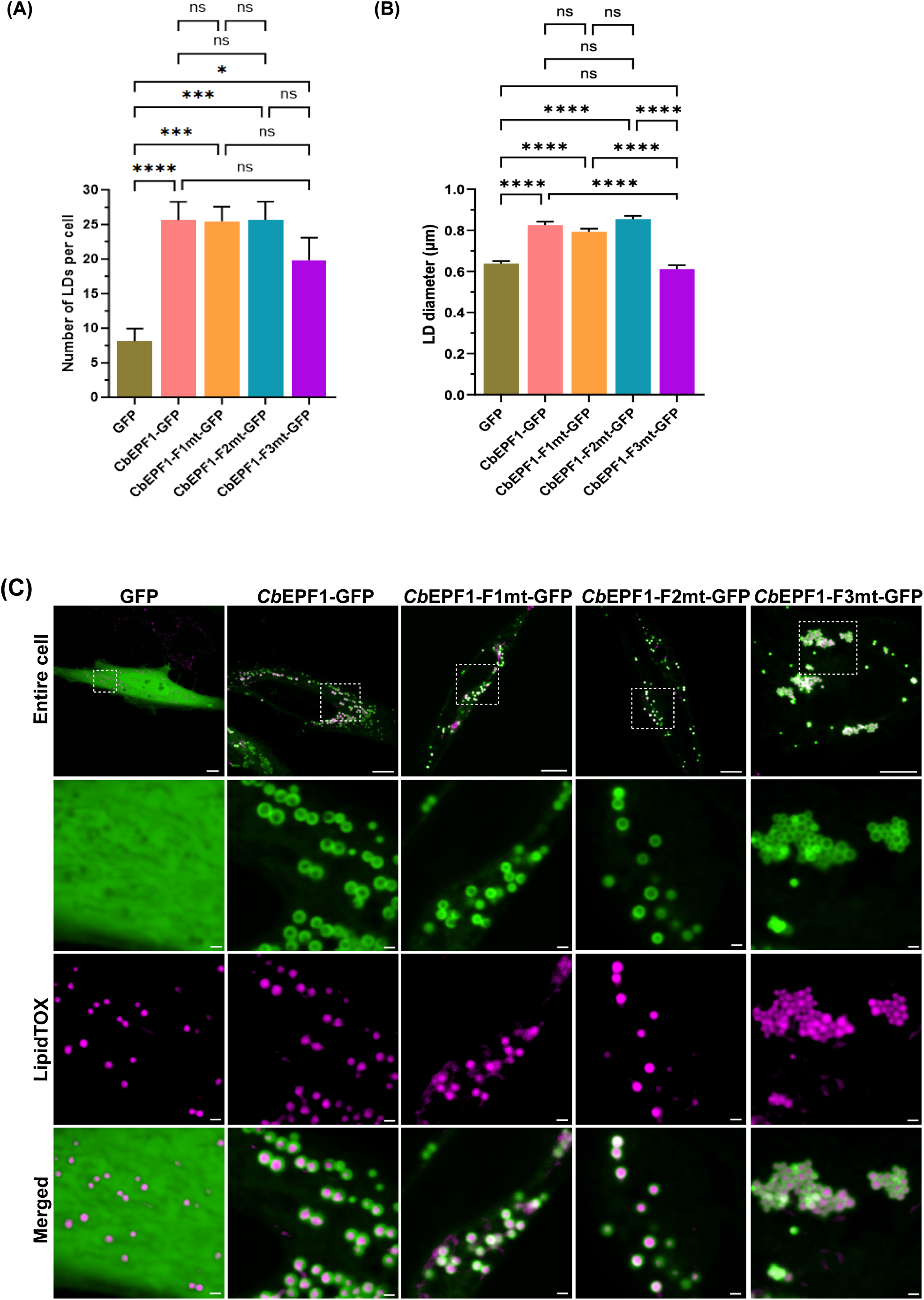
*Cb*EPF1 regulates host LD metabolism. (A) HeLa cells expressing *Cb*EPF1-GFP or *Cb*EPF1-FFAT mutant-GFP contain significantly high number of LDs compared to control cells. The number of LDs/cell in HeLa cells transiently expressing respective protein (GFP, n = 22 cells; *Cb*EPF1-GFP, n = 29 cells; *Cb*EPF1-F1mt-GFP, n = 25 cells; *Cb*EPF1-F2mt-GFP, n = 23 cells, *Cb*EPF1-F3mt-GFP, n = 25 cells) were quantified and shown as Mean ± SEM. One-way ANOVA with Tukey’s multiple comparisons test (ns, not significant; *, P < 0.05; **, P < 0.01; ***, P < 0.001; ****, P < 0.0001). (B) HeLa cells with *Cb*EPF1-GFP or *Cb*EPF1-F1mt-GFP or *Cb*EPF1-F2mt-GFP expression show larger LDs when induced with OA (100 μM) compared to GFP or *Cb*EPF1-F3mt-GFP expressing cells. The size of LDs in cells expressing respective protein (GFP, n = 90 LDs from 6 cells; *Cb*EPF1-GFP, n = 90 LDs from 8 cells; *Cb*EPF1-F1mt-GFP, n = 80 LDs from 6 cells; *Cb*EPF1-F2mt-GFP, n = 79 LDs from 7 cells, *Cb*EPF1-F3mt-GFP, n = 90 LDs from 7 cells) were quantified and shown as Mean ± SEM. One-way ANOVA with Tukey’s multiple comparisons test (ns, not significant; *, P < 0.05; **, P < 0.01; ***, P < 0.001; ****, P < 0.0001). (C) Representative images for LD size in HeLa cells expressing GFP or *Cb*EPF1-GFP or *Cb*EPF1-FFAT mutants, treated with OA (100 µM, overnight). Scale bars: 10 µm (overview) and 1 µm (magnified).

As *Cb*EPF1 induces ER-LD contacts, we sought to evaluate whether there is an impact on LD size upon OA treatment. We supplemented cells expressing *Cb*EPF1-GFP or *Cb*EPF1-GFP FFAT mutants with OA and measured the LD diameter. *Cb*EPF1-GFP expressing cells showed significantly larger LDs compared to cells expressing GFP. Expression of the single FFAT mutants, *Cb*EPF1-F1mt-GFP and *Cb*EPF1-F2mt-GFP, resulted in larger LDs compared to control GFP-expressing cells, but LD sizes relatively similar to *Cb*EPF1-GFP expressing cells. However, cells expressing the double FFAT mutant *Cb*EPF1-F3mt-GFP contained LDs similar in size to GFP control cells and significantly smaller LDs compared to *Cb*EPF1-GFP or *Cb*EPF1-F1mt-GFP or *Cb*EPF1-F2mt-GFP expressing cells (**Fig. 5B and 5C**). These results suggest that *Cb*EPF1 promotes the development of larger LDs in its FFAT motif dependent manner.

## DISCUSSION

Inter-organelle contact sites in eukaryotic cells serve as pivotal hubs for cellular homeostasis. Recent research has unveiled the strategies employed by pathogens to target these sites and regulate host cell metabolism (Dumoux and Hayward, 2016; Derre, 2017; Jiang et al, 2022). While viruses have been recognized for their manipulation of host inter-organelle contact sites, bacterial-mediated inter-organelle contact sites were previously limited to the ER and the BCV (Justis et al, 2017; Stoeck et al, 2018; Derre et al, 2019; Cook et al, 2022; Yue et al, 2023). Our study unveils the first example of a secreted bacterial effector protein that orchestrates inter-organelle contact sites beyond the confines of the BCV, thereby exerting control over host cellular lipid metabolism. Our results underscore that the FFAT motifs in *Cb*EPF1 play a central role in mediating inter-organelle contact sites between the host ER and LDs by interacting with VAPs in the host ER. Furthermore, these contact sites influence the growth of LDs, a critical aspect of lipid metabolism (**Fig. 6**).

**Figure 6:**
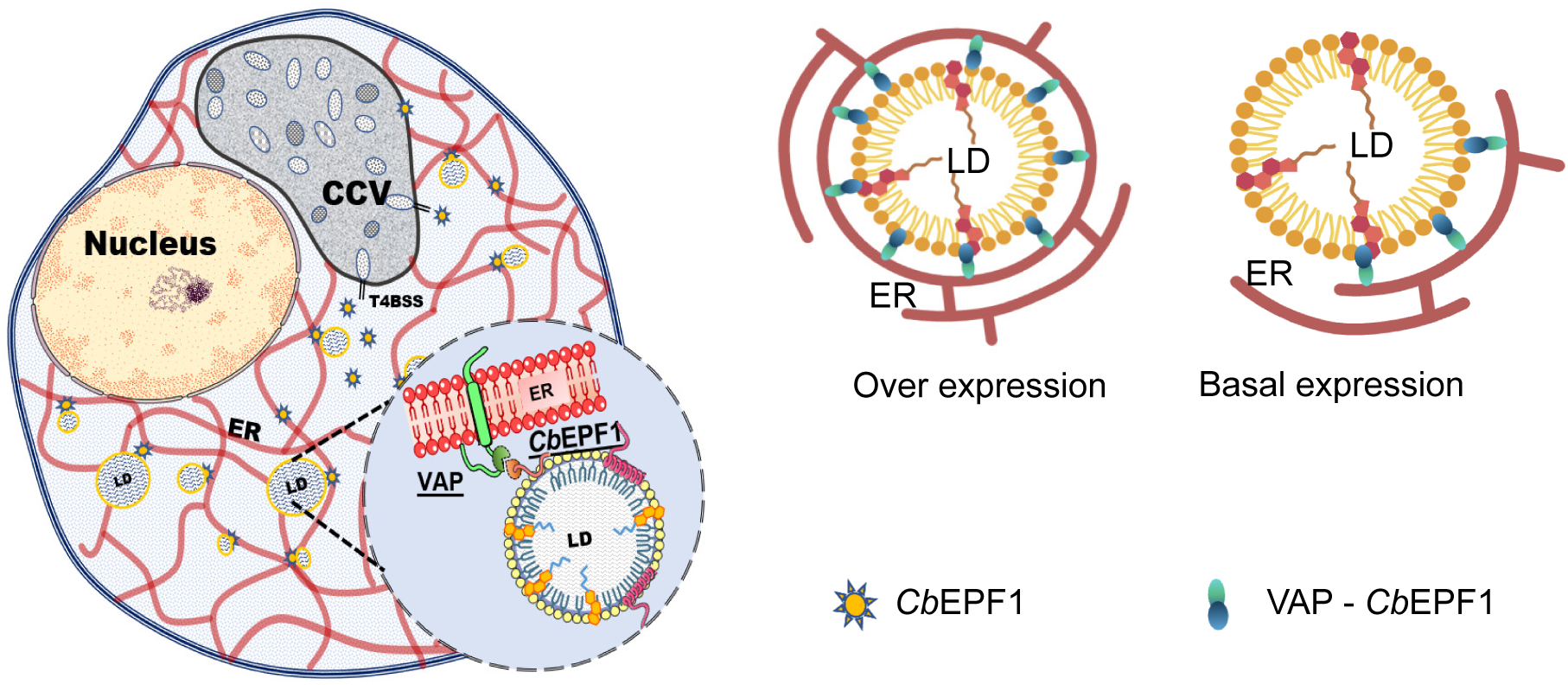
Model of the *Cb*EPF1-mediated inter-organelle contact sites between host ER and LDs to regulate host LD metabolism. Schematic representation of how *Cb*EPF1 establish ER-LD contacts in the *Coxiella* infected cells. *Cb*EPF1, localized on host LDs, interacts with VAPs in the ER to establish ER-LD contact sites (left side). The extent of *Cb*EPF1-mediated ER-LD contacts may be contingent on protein expression levels (right side). In overexpression conditions, *Cb*EPF1 promotes substantial ER wrapping around entire LDs. Conversely, in native cells with lower protein expression, *Cb*EPF1-mediated ER-LD contacts may be limited to specific short regions. The *Coxiella* Containing Vacuole (CCV) and type 4b secretion system (T4BSS) are depicted in the *Coxiella*-infected cell model.

Intriguingly, our findings reveal that *Cb*EPF1 exhibits functional parallels with eukaryotic proteins known to relocalize from ER to LD surface for LD development and growth. Notably, eukaryotic proteins such as GPAT4, DFCP1, LDAF1, and Rab18 relocalize from ER to LD surface, ultimately to promote LD growth (Wilfling et al, 2013; Xu et al, 2018; Chung et al, 2019; Li et al, 2019). While there is no sequence homology between *Cb*EPF1 and these eukaryotic proteins, our results suggest they serve similar functions. This functional similarity underscores the concept that bacterial effectors possess currently undiscovered domains or eukaryotic-like short linear motifs (SLiMs) as a mechanism to mimic host proteins and exert control over host cellular pathways. Significantly, mounting evidence supports the utilization of eukaryotic-like SLiMs by bacterial effectors as a strategy to hijack cellular machinery (Via et al, 2015; Samano-Sanchez and Gibson, 2020). The presence of FFAT motifs in *Cb*EPF1 serves as a compelling example of eukaryotic-like SLiMs in bacterial proteins and extends our understanding of these motifs and their adaptability in pathogenic contexts. The intriguing functional homology between *Cb*EPF1 and eukaryotic proteins raises questions about the precise mechanisms through which *Cb*EPF1 modulates host lipid metabolism. As the N-terminal region of *Cb*EPF1 is undefined, we cannot rule out the possibility of novel domains involved in synthesis of neutral lipids or sterol esters to promote LDs growth or expansion. While the presence of FFAT motifs suggests a *Cb*EPF1 role in lipid transport at ER-LD contacts, whether *Cb*EPF1 functions as a lipid transporter or as a molecular tether that recruits additional lipid transporters to the ER-LD contact sites has yet to be determined. The presence of two FFAT motifs in *Cb*EPF1 is an intriguing evolutionary adaptation that potentially enhances its interaction with VAPs in the ER, expanding the ER-LD contact area. Furthermore, the preferential interaction of the second *Cb*EPF1 FFAT motif with MOSPD2, a sole VAP that is known to localize on LD surfaces, raises possible complex functional role of *Cb*EPF1 on host LDs metabolism. *Cb*EPF1 underscores the pathogen’s remarkable capacity to manipulate host cell inter-organelle communication, emphasizing the need for further exploration of FFAT motif-mediated inter-organelle contacts in the context of pathogenic infections.

*C. burnetii* survives and replicates within a specialized *Coxiella*-containing vacuole (CCV) that is rich in sterols (Howe and Heinzen, 2016). However, our previous research demonstrated that the accumulation of excess cholesterol in the CCV membrane is detrimental to *C. burnetii* growth (Mulye et al, 2017). Consequently, *C. burnetii* has evolved multiple molecular strategies to mitigate the excessive cholesterol buildup on the CCV membrane, thereby avoiding toxicity (Justis et al, 2017; Clemente et al, 2022; Schuler et al, 2023). Notably, we previously described how *C. burnetii* hijacks the host sterol transporter ORP1L, which upon recruitment to the CCV membrane interacts with VAPs in the ER to establish inter-organelle contact sites between the ER and CCV. This interaction facilitates cholesterol efflux from the CCV to the ER (Justis et al, 2017 and Schuler et al, 2023). Our current study advances these findings by suggesting that *C. burnetii* possesses an active mechanism to redirect cholesterol away from the ER towards storage in LDs. *Cb*EPF1, which initially localizes to LD biogenesis sites in the ER and later translocates to LD surfaces, plays an active role in promoting LD growth. The functional similarities with eukaryotic proteins and its potential roles in ER-LD contacts and LD growth imply a multifaceted strategy employed by *C. burnetii* to manipulate host cellular pathways. Given that our present experiments relied on ectopic expression, future studies will benefit from the use of *C. burnetii Cb*EPF1 mutants as additional tools become available.

In conclusion, our results illuminate the multifaceted role of *Cb*EPF1 as a regulator of host LDs by intervening in host inter-organelle communication. Serving as a tether protein between the host ER and LDs through its FFAT motifs, *Cb*EPF1 showcases the ability of a bacterial effector protein to manipulate inter-organelle communication beyond BCVs. As we continue to delve into the intricacies of this interplay, future research should focus on deciphering the precise mechanisms through which *Cb*EPF1 modulates host lipid metabolism. Additionally, exploring the broader implications of *Cb*EPF1’s actions on the overall *C. burnetii* life cycle and pathogenesis could unveil new avenues for understanding the intricacies of host-pathogen interactions.

## MATERIALS AND METHODS

### Cell lines and Bacterial Strains

HeLa cells (ATCC CCL-2; ATCC, Manassas, VA, USA) were cultured in Roswell Park Memorial Institute (RPMI) 1640 medium (Corning, New York, NY, USA) supplemented with 10% fetal bovine serum (FBS; Atlanta Biologicals, Flowery Branch, GA, USA) and 2 mM l-alanyl-l-glutamine (Glutagro, Corning) at 37°C in a 5% CO_2_ atmosphere. HEK293T cells (ATCC CRL-3216) were cultured in Dulbecco’s modified Eagle medium (DMEM) with high glucose (Corning) with 10% FBS under similar conditions. All mammalian cell cultures were passaged every 3 days, and no cells older than 20 passages were used in experiments. Regular testing confirmed the absence of mycoplasma contamination in all cell lines. mCherry-expressing *C*. *burnetii* NMII (Beare *et al*., 2009) were grown for 4 days in ACCM-2, washed twice with phosphate buffered saline (PBS) and stored as previously described (Omsland et al, 2009). The *E. coli* Δ*cya* strain DHM1, generously provided by Dr. Amanda J. Brinkworth (University of Nebraska Medical Center), was propagated in LB broth (BD Difco, Franklin Lakes, NJ, USA) and plated on LB agar supplemented with appropriate antibiotics.

Cells were transfected using FuGENE6 (Promega, Madison, WI, USA) according to the manufacturer’s protocol. Where noted, cells were incubated with 30 or 100 μM oleic acid (Sigma-Aldrich, St Louis, MO, USA) complexed with fatty acid-free BSA for the indicated time to induce LD formation.

### Cloning and constructs

The *Cb*EPF1-GFP expression vector was constructed through PCR amplification using *C. burnetii* genomic DNA as the template with the following primers: forward primer 5’-atccctaccggtgatatgcctgataaaaccgacagtcttactatac-3’ and reverse primer 5’-ctttgctagccatccgcggaacttttccaatgtaacgaatccaagtcc-3’. The resulting PCR fragments were cloned into XhoI-linearized pT-Rex-C-GFP plasmid (Invitrogen, CA, USA) using In-Fusion HD cloning (Takara Bio, San Jose, CA, USA).

Mutants of *Cb*EPF1-GFP, including *Cb*EPF1-F1mt-GFP (Y232A), *Cb*EPF1-F2mt-GFP (F244A), and *Cb*EPF1-F3mt-GFP (YF/AA), were generated through site-directed mutagenesis using *Cb*EPF1-GFP as the template (Azenta Life Sciences, South Plainfield, NJ, USA).

The *Cb*EPF1-Wt expression vector was created using PCR amplification with *C. burnetii* genomic DNA as the template and the following primers: forward primer 5’-tgattacgccaagcttgatgcctgataaaaccgacagtcttactatac-3’ and reverse primer 5’-gcaggcatgcaagctaacttttccaatgtaacgaatccaagtcc-3’. The PCR fragments were ligated into the HindIII-linearized pKNT plasmid. Similarly, *Cb*EPF1-F1mt (Y232A), *Cb*EPF1-F2mt (F244A), and *Cb*EPF1-F3mt (YF/AA) variants were generated through site-directed mutagenesis (Azenta Life Sciences) using *Cb*EPF1-Wt as the template.

The VAPB-Wt expression vector was produced through PCR amplification using mCherry-VAPB (Addgene plasmid – 108126) as the template and the following primers: forward primer 5’-taccgagctcgaatttgatggcgaaggtggagcag-3’ and reverse primer 5’-ttatatcgatgaattctacaaggcaatcttcccaataattacacca-3’. The PCR fragments were ligated into the EcoRI-linearized pUT18C plasmid. The VAPB-MSPmt expression vector was generated using pEGFPC1-hVAP-B KD/MD (addgene plasmid – 104450) as the template, harboring mutations in the MSP domain (K87D and M89D), with the same primers and cloning strategy as for VAPB-Wt. mCherry-VAPB (Addgene – 108126), pEGFPC1-hVAP-B KD/MD (Addgene – 104450), pEFIRES-P-mTagBFP-KDEL (Addgene – 87163) were purchased from Addgene. All plasmid constructs were validated through Sanger didexoy sequencing (ACGT, Inc., Wheeling, IL, USA) and Oxford Nanopore Technology (Plasmidsaurus, Eugene, OR, USA)

### Microscopy

All experiments were conducted by live cell imaging on a Nikon Eclipse Ti2 spinning disc microscope within an environmental chamber (5% CO_2_ and 37°C). Cells transfected with plasmids were cultured on ibidi slides (ibidi USA, Inc., Fitchburg, WI, USA), and imaged 48 hours post-transfection. For experiments involving oleic acid treatment, growth media was supplemented with 100 µM oleic acid for 16 hours before imaging. To observe the association of *Cb*EPF1 with LD biogenesis, time-lapse movies were acquired at 1-minute intervals for 20 minutes. Data quantification and image processing were carried out using the Nikon NIS-elements software. The Alpha-blending algorithm in NIS elements software was used for 3D rendering.

### Bacterial Adenylate Cyclase-based Two-Hybrid assay (BACTH)

The BACTH assay was performed as previously described (Ouellette et al, 2017). Briefly, the *E. coli* Δ*cya* strain DHM1 was co-transformed with pKNT25 and pUT18C derivatives. Following transformation, the bacteria were plated on LB agar supplemented with 5-bromo-4-chloro-3-indolyl-β-d-galactopyranoside (X-gal, 40 µg ml−1), IPTG (1 mM), Carbenicillin (100 μg ml−1), and Kanamycin (50 μg ml−1) and incubated for 48 hours at 30°C. Blue color of the colonies indicate a positive interaction between the analyzed proteins, while white color denotes the absence of complementation.

### Immunoprecipitation

HEK293T cells were seeded at a density of 1×10^5^ cells per well in a 6-well plate and transfected with GFP or *Cb*EPF1-GFP or *Cb*EPF1-F1mt-GFP or *Cb*EPF1-F2mt-GFP or *Cb*EPF1-F3mt-GFP, using Fugene6 transfection reagent (Promega). After 24 hours of transfection, cells were treated with 100 µM oleic acid for 16 hours. Subsequently, cells were collected and lysed on ice for 30 minutes using an immunoprecipitation (IP) buffer (15 mM Tris pH 7.4, 150 mM NaCl, 1% Triton X-100, 1x protease inhibitor cocktail (CST, Danvers, MA, USA). The supernatant obtained after centrifugation at 13,000 xg, 4°C for 10 minutes was incubated with GFP-Trap magnetic beads (Bulldog Bio, Inc., Portsmouth, NH, USA) at 4°C overnight. Following the incubation, the beads were washed with the IP buffer and bound proteins eluted and analyzed using SDS– polyacrylamide gel electrophoresis (SDS-PAGE) followed by immunoblotting with appropriate antibodies.

### Western blot analysis

Proteins were separated on 4-20% Mini-PROTEAN TGX stain-free protein gels (Bio-Rad, Hercules, CA, USA) and transferred to a nitrocellulose membrane. The membrane was blocked with 5% skimmed milk in TBS-T for 1 hour at room temperature. The membranes were washed with 1x TBST, incubated with primary antibodies at 1:1000 dilutions in 1% BSA-TBST overnight at 4°C, and then washed with 1x TBST prior to incubation with secondary antibodies coupled to horseradish peroxidase. Signals were detected using an enhanced chemiluminescence system (Thermo Fisher Scientific, Inc., Waltham, MA, USA) and visualized with Azure 600 western blot imager (Azure biosystems, Dublin, CA, USA). The primary antibodies used were anti-GFP (Sigma Aldrich, G6539), anti-MOSPD2 (Sigma Aldrich, ZRB1046), and anti-VAPB (Proteintech, Rosemont, IL, USA, 14477-1-AP).

### LD measurements

Lipi-Blue (Dojindo Molecular Technologies, Rockville, MD, USA) and HCS LipidTOX Red neutral lipid stain (Invitrogen) were used for LD staining. LD measurements were performed as previously described (Li et al, 2019). LD numbers / cell were manually quantified in the absence of oleic acid treatment using Nikon NIS Elements AR software. LD diameter was measured after treatment with 100 μM oleic acid using Nikon NIS Elements AR software. The cells or images for LD analysis were chosen randomly.

### Statistical analysis

The statistical parameters, including n, mean and SEM, are reported in the corresponding figure legends. Statistical analysis was performed in Prism (GraphPad, San Diego, CA, USA). Statistical significance was calculated by one-way ANOVA analysis. P-values <0.05, <0.01, <0.001, and <0.0001 are identified with *, **, ***, and ****, respectively. A P value > 0.05 was considered non-significant (ns).

## Acknowledgements

We thank Dr. Micah Schott for critical comments on the manuscript. We thank Gilk lab members for helpful discussions and critical feedback. We thank Dr. Amanda Brinkworth for providing plasmids for BACTH assay. This work was supported National Institutes of Health (AI139176 to S.D.G.) and the American Heart Association (906475 to R.K.A.). Schematic model was created using Servier Medical Art (http://smart.servier.com) and Biorender software (http://Biorender.com).

## Author contributions

RKA and SDG designed the experiments. RKA and AS performed experiments. RKA and SDG wrote the manuscript.

The authors declare no competing financial interests.

## MATERIALS & CORRESPONDENCE

Further information and requests for reagents may be directed to and will be fulfilled by Stacey Gilk

## DATA AVAILABILITY

The data generated and analyzed in this study are available from the corresponding author upon request.

---

**Supplemental Figure 1:**
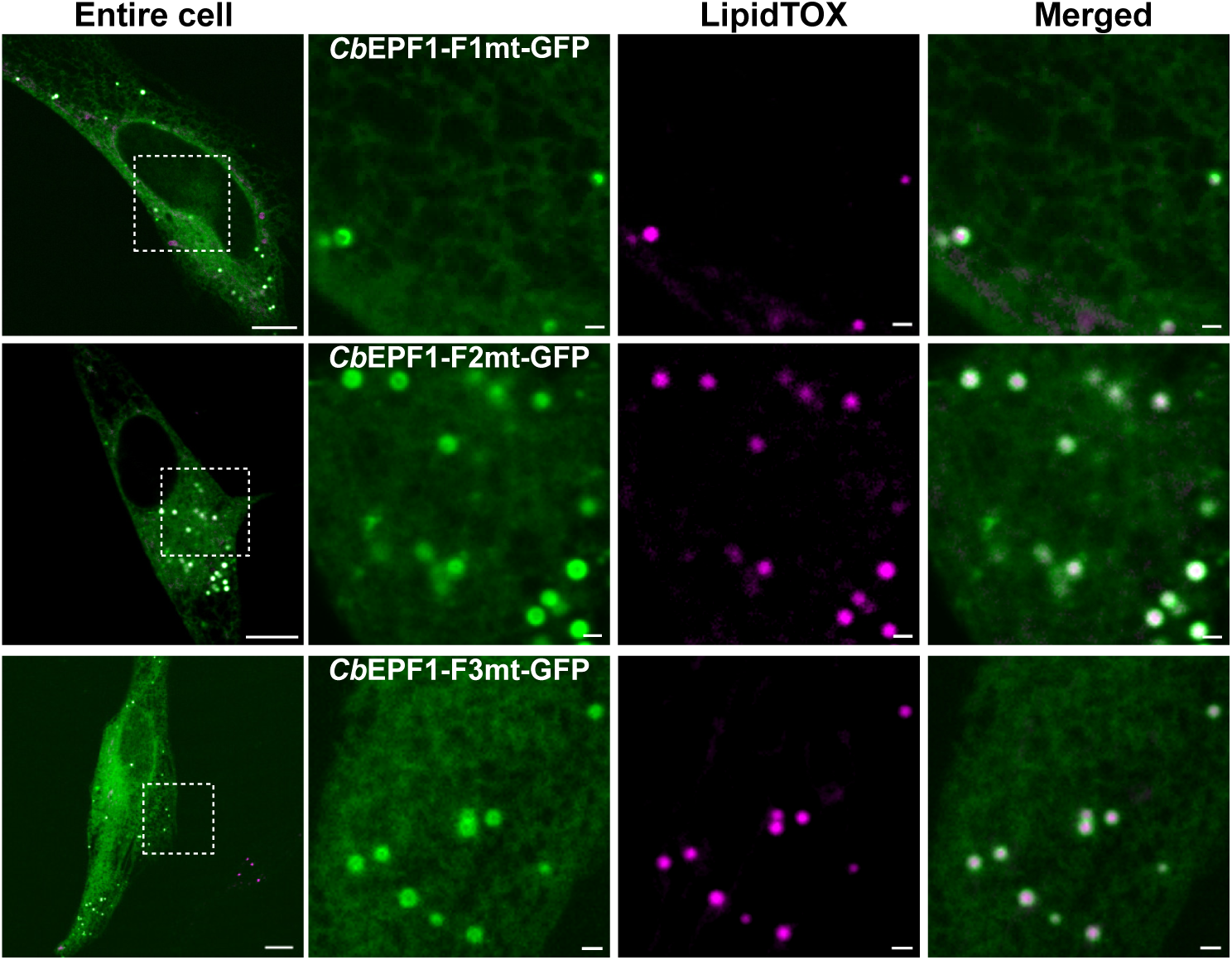
The ectopically expressed *Cb*EPF1-F1mt-GFP or *Cb*EPF1-F2mt-GFP or *Cb*EPF1-F3mt-GFP localized to ER and LDs in HeLa cells. The *Cb*EPF1-FFAT mutant-GFP proteins showed a similar localization pattern to *Cb*EPF1-GFP (Fig. 1B). The LDs were labeled with LipidTOX-Red. Scale bars: 10 µm (overview) and 1 µm (magnified).

## Notes

### Competing Interest Statement

The authors have declared no competing interest.

